# Reappraising highly arousing negative content recruits the ventromedial prefrontal cortex in older adults

**DOI:** 10.64898/2026.11.24.768795

**Authors:** L. Ceravolo, G. Moy, C. Simon, A. Vazquez, D. Framorando, G. Gold, F. Lazeyras, P. Vuilleumier, P. Giannakopoulos, C. Delaloye

**Author notes:** Correspondence: Leonardo Ceravolo, PhD University of Geneva & Swiss Center for Affective Sciences Biotech Campus 9, Chemin des Mines CH-1202 Geneva, Switzerland.

## Abstract

Coping with negative events occurs on a daily basis during adulthood, giving importance to the study of emotional regulation strategies and their underlying brain mechanisms. While passively processing a negative scene (“watching”) may sometimes lead to acceptable emotional regulation, an extensive body of literature highlighted the improved efficiency of an active, cognitive evaluation of such events (“reappraising”), emphasizing interindividual differences related to age groups, most notably between younger and older adults. In the present study, we investigated whether reappraising as opposed to watching pictures depicting negative events would lead to a decrease in reports of negative feeling in a group of older adults, and whether this decrease would rely on activity in brain regions typically found in this literature for adult populations—namely, the amygdala and the ventromedial prefrontal cortex. Our behavioral results revealed the expected decrease of negative feeling for the reappraisal condition and reappraising (relative to watching) negative pictures was associated with enhanced activity in the left supramarginal and middle temporal gyri. Additional analyses revealed a strong impact of the arousing value of the pictures, emphasizing increased brain activity in the ventromedial prefrontal cortex in relation to increasing arousal of the images that were reappraised rather than simply watched. Exploratory seed-to-seed functional connectivity analyses further indicated that reappraising, relative to watching, was associated with stronger coupling of the anterior and posterior cingulate cortices with right lateral prefrontal regions, and of the medial frontal cortex with the right cerebellum. Our results confirm that the ability to effectively reappraise (as opposed to watch) the content of negative events is preserved in older adults. However, activity in the ventromedial prefrontal cortex seems to be specifically enhanced for highly arousing stimuli, a result that interestingly questions the ability of static pictures to elicit a strong emotional response in older adults.

## 1 Introduction

Emotion perception occurs when we encounter emotional events in our everyday life, which is a rather frequent phenomenon. In some cases, the mere exposition to an emotional event may trigger some mechanisms known to help the individual to cope with this specific situation. Among the processes that take place after or even during the event, relevant appraisals (Scherer, 1984) are usually complemented by emotional regulation strategies (Gross, 1998, 1999) such as reappraisal for instance (Thiruchselvam et al., 2011). Reappraising an emotional event may be the most reliable strategy (since the individual is active in the process), and it is done by cognitively reassessing the event in order to extract meaning from it. It also implies to find an interpretation that suits the perceiver in particular and decreases the impact of the event for him (Gross, 1998). In opposition with reappraisal, suppression involves the individual to block the emotional input while the last and most passive way of processing a relevant event is to simply watch it, without the use of any strategy or diminishing factor whatsoever.

As one can infer from the literature mentioned above, a failure to use the most adapted strategy for emotion regulation (or to use one at all) can have consequences regarding the subjective feeling of the individual, especially when encountering a negative event. In fact, several psychopathologies such as depression were shown to crucially depend on regulation strategies frequently employed by an individual when facing negative situations (Gross and Muñoz, 1995). Cognitive emotion regulation was shown to be a key factor in major depression (Erk et al., 2010) and the use of spontaneous (as opposed to instructed) suppression over reappraisal was shown to happen more often in recovered-depressed patients as compared to healthy controls (Ehring et al., 2010). Reappraisal was interestingly shown to rely on the anterior cingulate and prefrontal cortex (Ochsner et al., 2012), brain regions known to also play a key role in depression (Ebert and Ebmeier, 1996; Johnstone et al., 2007) in addition to the amygdala and other subcortical structures (Wager et al., 2008). Therefore, emotion regulation and depression seem to be linked both at the cognitive and neural levels, making the study of emotion regulation mechanisms a crucial topic in helping improve our understanding and treatment of depression, be it for younger adults or older adults. Indeed, extensive literature highlights the existence and whereabouts of depression in the general population, but late life appears to be a critical time-window for depression to take place (Blazer, 2003).

Nuancing these findings, an expanding literature showed that arousing material such as negative pictures would not affect younger and older adults in a similar manner. More specifically, the tendency to automatically focus one’s attention on negative material (negativity bias) was shown to be more predominant in younger than older adults (Wood and Kisley, 2006). This observation could reflect age-related differences in perceptual and attentional mechanisms but it should also be noted that the task itself has a tremendous impact on subjective feeling and physiology (arousal), and older adults seem to be especially sensitive to this aspect (for a review, see Lau et al., 2001).

For the reasons stressed above, the present study aimed at a clearer and specific understanding of cognitive emotional regulation mechanisms and their underlying brain substrates in a population of healthy older adults (cognitive assessment and neuropsychological testing were also performed). To do so, we used a simple task in which the participants were presented with emotional (negative) and neutral pictures. Following picture presentation, their task was to assess their emotional feeling during picture perception across instructed cognitive (reappraisal) and perceptual (watch) conditions. During the whole task, BOLD (Blood oxygenation level-dependent) functional magnetic resonance imaging data were acquired continuously. Our hypotheses were the following: i) because reappraising as opposed to simply watching visual content is supposed to lead to a better emotional feeling, we expected higher negative feelings reported for watch than reappraise conditions; ii) we also expected higher negative emotions felt during negative as compared to neutral content presentation; iii) enhanced BOLD signal was also expected in the anterior part of the cingulate cortex and in the ventromedial prefrontal cortex (vmPFC) as well for reappraisal as compared to watch conditions; iv) highly arousing stimuli were expected to influence BOLD signal in the vmPFC and amygdala, especially for the reappraise and watch condition, respectively; v) finally, at the network level, we expected reappraisal relative to watching to strengthen functional coupling between cingulate and lateral prefrontal control regions, as well as between these regions and areas involved in the appraisal of emotional content.

## 2 Methods

### 2.1 Sample size and power estimation

It was shown that 24 participants for an fMRI study accounted for at least 80% of the variance at the voxel level, as estimated for a conservative [0.01<α<0.05] alpha (Desmond and Glover, 2002). Sample size was therefore calculated using G*Power 3 software (Faul et al., 2007) with standard values for the architecture of our study (power of 0.95 and alpha of 0.05) given a sample of 24 participants and only two conditions. With that sample size, power estimation reached 86.22%. We hence recruited 24 participants matching our inclusion criteria.

### 2.2 Participants and inclusion criteria

Twenty-four right-handed, healthy, native or highly proficient French-speaking participants (15 male, 9 female, mean age 73.4 years, SD 6.4, range 63-84) were included in the study. All participants had normal hearing, normal or corrected to normal vision and no history of psychiatric or neurologic disease. They all gave written informed consent for their participation in accordance with ethical and data security guidelines of the University of Geneva and of Geneva University Hospital. The study was approved by the cantonal research ethics committee of Geneva (Commission Cantonale d’Éthique de la Recherche, CCER) and was conducted according to the Declaration of Helsinki.

In order to avoid any confounds or biases linked to depression in our sample, all participants were assessed using the 15-item Geriatric Depression Scale (Yesavage et al., 1983). All participants included in the present study had a score below 5 (mean 1.4; SD 1.3; range 0-4). The semi-structured Mini International Neuropsychiatric Interview (Lecrubier et al., 1997) was also administrated to the participants by a trained clinical psychologist in order to exclude any past or current psychiatric disease. No participant had any known psychiatric disease. Eventually, the Mattis Dementia Rating Scale (Mattis, 1976) was administrated to the participants by a trained neuropsychologist in order to control for any cognitive impairment. All participants had a score within the normal range (mean 138.5; SD 3.6; range 131-144).

### 2.3 Neuroimaging task stimuli

Stimuli were selected from the International Affective Picture System (IAPS) and included pictures representing scenes involving either neutral or negative valence (sadness, disgust, anger). These pictures underwent a normative procedure regarding their valence, arousal and dominance (Lang et al., 2008) and are thus highly validated. We however had them evaluated by an independent sample that was age-matched to our fMRI participants (N=22; 12 male, 10 female, mean age 67.8, SD 5.9, range 60-82). Stimuli arousal and valence were evaluated by the independent sample (Negative images: mean arousal=7.50, SD=0.45, mean valence=2.24, SD=0.54; Neutral images: mean arousal=4.76, SD=0.38, mean valence=4.75, SD=0.44). They were presented in the fMRI through a rear-mounted projector with a mirror attached to the upper part of the 12-channel MRI head-coil.

### 2.4 Task and experimental design

Participants had to report whether they felt negative emotions during the presentation of the picture by pressing one of the four available keys (MR-compatible Response box: Current Designs Inc., Philadelphia, PA, USA) (left key: no negative emotions felt; middle left key: mild negative emotion felt; middle right key: mild to high negative emotion felt; right key: high to extreme negative emotion felt; responses were coded from 1 to 4, respectively). Two different conditions were presented: the watch condition and the reappraise condition. In the watch condition, participants had to look at the picture and evaluate it, whereas, in the reappraise condition, they were asked to try to decrease their subjective feeling by finding a cognitive strategy. The instructions related to the reappraise condition were explained in detail before the task by giving examples of various strategies that could be used (for instance: “You could try to find positive aspects to the scene”, “You could imagine that the image is a fake”). Besides, an extended training session was administrated to the participants before the task.

The task was administered in two functional runs of approximately 12.5 minutes each. Each run comprised 12 mini-blocks of 4 pictures (6 Watch and 6 Reappraise mini-blocks), presented in alternating series of three mini-blocks of the same condition (three Reappraise followed by three Watch mini-blocks, repeated twice). A given condition remained constant during each mini-block. Each mini-block contained three negative pictures and one neutral picture, the neutral picture appearing in the second, third or fourth position. Each run thus included 18 negative and 6 neutral trials per condition, and the same 48 pictures (36 negative, 12 neutral) were presented in both runs, each picture always being associated with the same condition. Stimulus lists were counterbalanced across participants. Each mini-block was announced by a countdown screen (approximately 8 seconds), after which the instruction corresponding to the block (Watch or Reappraise) was displayed for 5 seconds to allow the participants to get ready (see Figure 1). Then, the first picture was presented for 8 seconds, followed by a response slide of 3.5 seconds (during which participants had to give their answer) and a blank screen of 1 second. Then, the second picture was presented followed by a response slide and a blank within the same timeframe and so on. The instruction remained written below each picture and the four possible responses were also written on the response slide to allow the participants to go through the task without having to remember the instruction and possible responses. In the second run, a short caption was additionally displayed below each picture: in Reappraise trials, it suggested a possible reappraisal of the scene (e.g., “It is a scene from a movie”), whereas in Watch trials, it described the content of the scene (e.g., “Tornado in the distance”).

**Figure 1.**
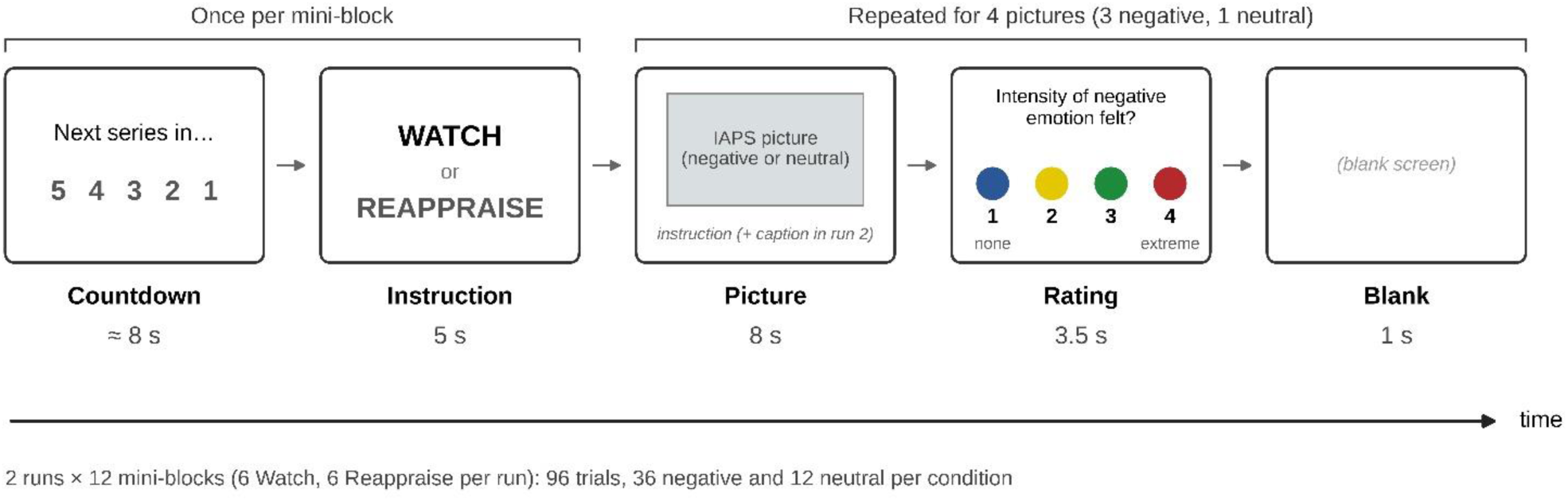
Experimental design. Each mini-block was announced by a countdown screen (approximately 8 s), followed by the instruction (Watch or Reappraise; 5 s) and four trials, each consisting of an IAPS picture presented with an instruction reminder (8 s; in the second run, a short caption suggesting a reappraisal or describing the scene was also displayed), a rating screen on which participants reported the intensity of the negative emotion felt on a 4-point scale (1 = no negative emotion to 4 = high to extreme negative emotion; 3.5 s), and a blank screen (1 s). Each mini-block contained three negative pictures and one neutral picture. The task comprised two runs of 12 mini-blocks (6 Watch, 6 Reappraise per run), for a total of 96 trials (36 negative and 12 neutral trials per condition).

There were hence 48 trials in total for each condition (48 Watch/48 Reappraise) for a grand total of 96 trials across conditions. For each condition, 36 of the trials had a negative valence, 12 had a neutral valence.

### 2.5 Behavioral data analysis

Following normality estimation, data were analysed by using a 2 (Condition) x 2 (Emotion) repeated measures analysis of variance (ANOVA) with Statistica 12 software (StatSoft Inc., Tulsa, OK, USA).

Ratings (1–4) were first averaged across the two runs for each participant and each Condition × Emotion cell. Significant effects were followed up with two-tailed paired t-tests. Effect sizes are reported as partial eta squared (ηp²) for ANOVA effects and Cohen’s dz for paired t-tests. Because ratings for neutral images were close to the floor of the scale, follow-up comparisons were additionally verified with Wilcoxon signed-rank tests.

### 2.6 Neuroimaging data acquisition

Structural and functional brain imaging data were acquired by using a 3T scanner (Siemens Trio, Erlangen, Germany) with a 12-channel coil. A magnetization prepared rapid acquisition gradient echo sequence was used to acquire high-resolution (1 x 1 x 1 mm3) T1-weighted structural images (TR = 1,900 ms, TE = 2.27 ms, TI = 900 ms). Functional images were acquired by using a multislice echo planar imaging sequence (36 transversal slices in descending order, slice thickness 3.2 mm, TR = 2,100 ms, TE = 30 ms, field of view = 205 x 205 mm2, 64 x 64 matrix, flip angle = 90°, bandwidth 1562 Hz/Px).

### 2.7 Neuroimaging data analysis

Functional images were analysed with Statistical Parametric Mapping software (SPM12, Wellcome Trust Centre for Neuroimaging, London, UK). Preprocessing included realignment to the first volume of the time series, slice timing, normalization into the Montreal Neurological Institute (MNI) (Collins et al., 1994) space was improved by using the DARTEL toolbox (Ashburner, 2007) and spatial smoothing was computed with an isotropic Gaussian filter of 8 mm full width at half maximum. To remove low frequency components, we used a high-pass filter with a cut-off frequency of 128s. Anatomical locations were defined with a standardized coordinate database (Talairach Client, http://www.talairach.org/client.html) by transforming MNI coordinates to match the Talairach space (and then transforming it back into MNI).

We used one general linear model at the first level (subject-level analysis). In this model, each regressor was modeled by using a stick function that included stimulus duration as well and was then convolved with the hemodynamic response function. Events were time-locked to the onset of the image stimuli. Separate regressors were created for both conditions and their valence (Watch negative/neutral, Reappraise negative/neutral). Image arousal obtained through evaluation by an independent, age-matched sample was included here as a parametric modulator on a trial-by-trial basis. Parametric modulators were serially orthogonalized with respect to their condition regressor (SPM12 default). Additionally, six motion parameters were included as regressors of no interest to account for movement in the data, and no participant moved more than 3mm (translation) or 3° (rotation).

The condition regressors and the parametric modulators were used to compute linear contrasts for each participant and were taken to two second-level (group-level analysis), flexible factorial analyses. The first second-level analysis (Model 1) was performed using the Condition as within-subject factor, with a 2(Watch, Reappraise) factorial structure. The Condition factor was included in order to test differences between our conditions, namely Watch>Reappraise and the inverse contrast. Since each condition included 36 negative and only 12 neutral stimuli regarding valence, only negative stimuli were included at the second-level of analysis for model 1. This decision was also taken in order to specifically test the mechanisms underlying the cognitive reappraisal of negative events, which was the general goal of the present study.

The other second-level analysis (Model 2) was performed on the first-level parametric modulator estimates, that is, on the slopes relating the trial-by-trial arousal value of the images to the BOLD signal in each condition. Arousal slope was entered as a within-subject factor with a 2(Watch arousal, Reappraise arousal) factorial structure. This model thus tested whether the relationship between image arousal and brain activity differed between conditions: a Reappraise arousal > Watch arousal effect indicates that the BOLD signal increased more steeply with image arousal during reappraisal than during watching (and conversely for Watch arousal > Reappraise arousal), rather than a difference in mean activity between conditions. This factor was also included in order to take into account the arousing value (indirectly linked to the emotional value) of the stimuli, especially since we could not directly compare them regarding valence due to an insufficient number of trials. For both models (Model 1 & 2), flexible factorial second-level analyses were computed with one main effect of Subject (mandatory in SPM12 to account for inter-individual variance), one main effect of Condition (Model 1) and one main effect of Arousal slope (Model 2). Both models assumed that participants were independent while Condition (Model 1) and Arousal slope (Model 2) were not. Finally, variance estimation was set to unequal in both models for all factors (Subject factor, Condition and Arousal slope factors) because homoscedasticity cannot be evaluated and data cannot be transformed.

All activations are reported at an intensity threshold of p < 0.005 (uncorrected) and a cluster extent threshold of k > 86 voxels, equivalent to a Family-Wise Error correction for multiple comparison of p < .05 at the cluster level. This threshold was based on the final smoothing of the data and determined with the *3dClustSim* function in AFNI (Cox, 1996) software (http://afni.nimh.nih.gov/afni). The *3dClustSim* function uses a non-parametric method with 10,000 iterations to estimate the necessary cluster extent thresholding for side-to-side voxels (NN-2 option was used here). *3dClustSim* reports a cluster extent threshold for each specified statistical *p* value and assumes that neighboring voxels are part of a similar functional response pattern, rather than taking each voxel as a completely different and independent measure as implied by the family-wise error correction at the voxel level. Statistically significant activation maps were then displayed on an age-specific (Template mean age: 65 years old; Rorden et al., 2012) anatomical mean MRI template, which is part of the clinical-toolbox for SPM (Clinical-toolbox: http://www.nitrc.org/frs/?group_id=881). This procedure was used to better match the anatomical structure of our sample with a validated, mean anatomical image and the above-mentioned template was the closest to our sample’s age range and had the highest quality and resolution.

Seed-to-seed (ROI-to-ROI) functional connectivity (FC) analyses were performed with the CONN toolbox (Whitfield-Gabrieli and Nieto-Castanon, 2012) version 18.b implemented in Matlab 9.0 (The MathWorks, Inc., Natick, MA, USA), using the preprocessed (realigned, slice-time corrected, DARTEL-normalized and smoothed) functional images described above. Regions of interest (ROIs) were defined with CONN’s default atlas, which comprises 132 ROIs: 91 cortical and 15 subcortical regions from the Harvard-Oxford atlas (Frazier et al., 2005; Desikan et al., 2006; Makris et al., 2006; Goldstein et al., 2007) and 26 cerebellar and vermal regions from the Automated Anatomical Labelling atlas (Tzourio-Mazoyer et al., 2002). For each participant, the BOLD time series of each ROI was obtained by averaging the signal across all voxels of that ROI from the smoothed functional data.

Spurious sources of noise were removed with CONN’s default denoising pipeline, which combines linear regression of confounding effects with temporal band-pass filtering. Confounds included five principal components each extracted from white matter and cerebrospinal fluid masks (anatomical component-based noise correction, aCompCor; Behzadi et al., 2007), 12 realignment parameters (the six motion parameters of each of the two runs) and the main effects of each task condition convolved with the canonical hemodynamic response function together with their first-order derivatives, so that connectivity estimates would not merely reflect shared task-evoked co-activation. The residual BOLD time series were then linearly detrended and band-pass filtered (0.008 < f < 0.09 Hz).

At the first level, condition-specific connectivity was estimated with generalized psychophysiological interaction (gPPI) models (McLaren et al., 2012), using bivariate correlation as the connectivity measure. For each seed–target pair of ROIs, the gPPI model included the seed ROI time series (physiological term), the condition regressors convolved with the canonical hemodynamic response function (psychological terms), and the product of each condition regressor with the seed time series (interaction terms). The interaction terms quantify how strongly the seed and target time series are coupled during each condition, over and above task-evoked co-activation, and were taken as condition-specific connectivity estimates. Because gPPI estimates are directional (the seed and target roles are not interchangeable), each connection is reported below as seed–target. Each condition was entered separately as a simple effect, that is, one regressor per condition, valence and run (Watch negative, Watch neutral, Reappraise negative and Reappraise neutral, for each of the two runs).

At the group level, gPPI connectivity estimates were compared between conditions with a within-subject contrast (Reappraise > Watch; paired t-tests, N = 24). Statistical inference followed CONN’s seed-level approach: for each seed ROI, an omnibus F-test first assessed whether its overall connectivity profile differed between conditions, and individual connections were thresholded at p < 0.05 false discovery rate (FDR; Benjamini and Hochberg, 1995) corrected for the 131 connections of that seed (seed-level correction). Because this procedure controls the false discovery rate within each seed rather than across all 8,646 connections of the network, these connectivity results are reported as exploratory.

## 3 Results

### 3.1 Behavioral data

To compare our two conditions, namely Watching or Reappraising the content of images presented in mini-blocks, we computed a 2 (Condition: Watch, Reappraise) × 2 (Emotion: negative, neutral) repeated-measures ANOVA on felt emotional intensity ratings (higher scores indicating a more intense negative emotion). As expected, we observed significant main effects of Condition (F(1,23) = 41.59, p < .001, η_p_² = .64) and Emotion (F(1,23) = 250.98, p < .001, η_p_² = .92), as well as a significant Condition × Emotion interaction (F(1,23) = 20.58, p < .001, η_p_² = .47; Figure 2).

**Figure 2.**
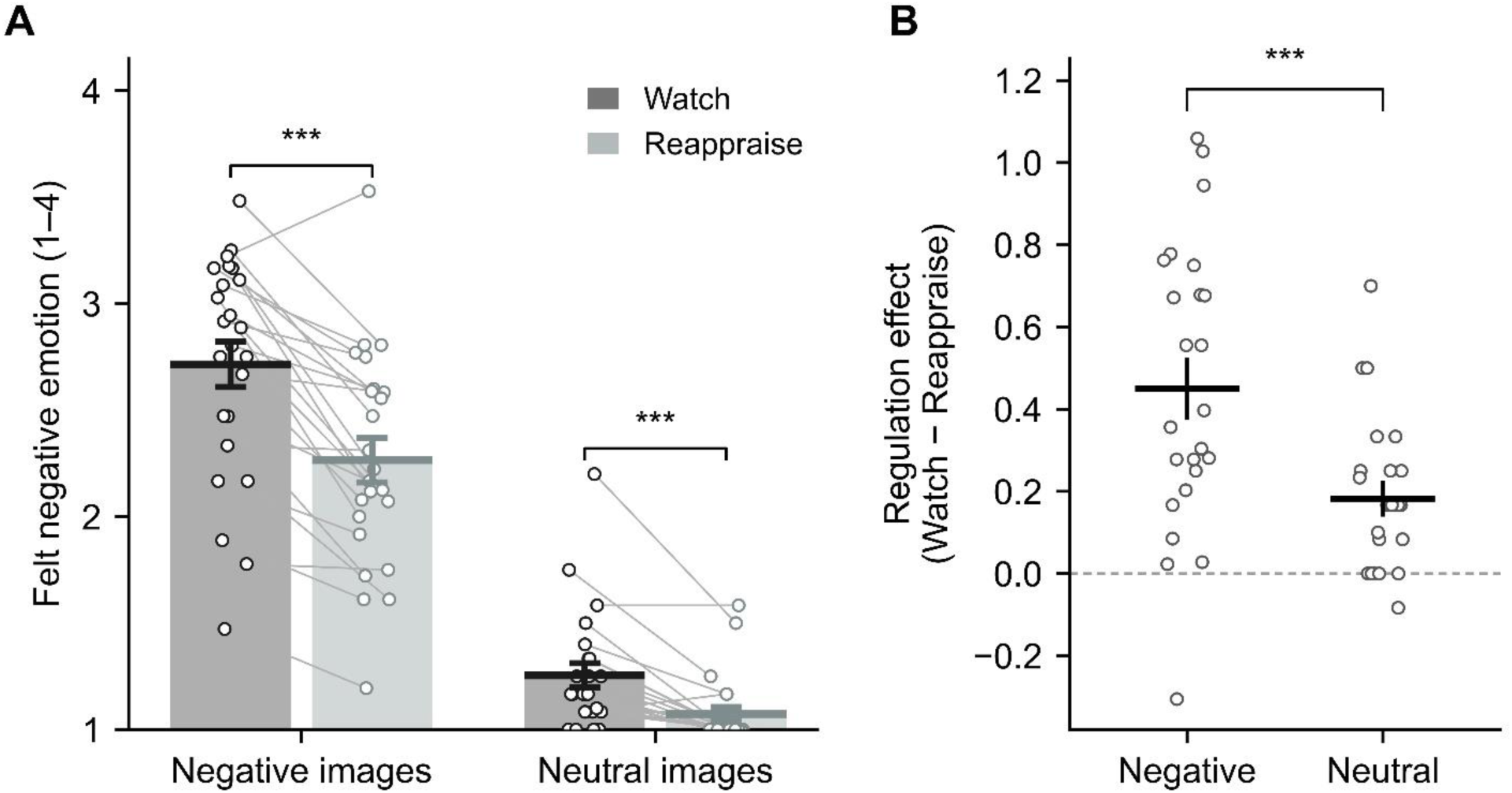
Felt emotional intensity. (A) Mean ratings (± SEM) for Watch and Reappraise trials, separately for negative and neutral images; dots and grey lines depict individual participants (N = 24). (B) Regulation effect (Watch − Reappraise) for negative and neutral images; horizontal bars show the mean ± SEM, dots depict individual participants. ***p < .001, paired t-tests.

The main effect of Condition was explained by felt emotional intensity being scored higher for Watch (M = 1.98, SD = 0.36) than for Reappraise trials (M = 1.67, SD = 0.31; t(23) = 6.45, p < .001, dz = 1.32). For the Emotion factor, scores were higher for negative (M = 2.49, SD = 0.49) than for neutral images (M = 1.16, SD = 0.21; t(23) = 15.84, p < .001, dz = 3.23), showing that our participants were moved in a stronger manner by negative relative to neutral content pictures.

Follow-up comparisons of the Condition × Emotion interaction showed that reappraising, compared to watching, decreased felt emotional intensity both for negative images (Watch: M = 2.71, SD = 0.52; Reappraise: M = 2.26, SD = 0.51; t(23) = 6.34, p < .001, dz = 1.29) and for neutral images (Watch: M = 1.25, SD = 0.28; Reappraise: M = 1.07, SD = 0.16; t(23) = 4.69, p < .001, dz = 0.96; Wilcoxon signed-rank test, p < .001; Figure 2A). Crucially, this regulation effect (Watch − Reappraise) was more than twice as large for negative (M = 0.45, SD = 0.35) as for neutral images (M = 0.18, SD = 0.19; Figure 2B), which accounts for the interaction.

### 3.2 Neuroimaging data

#### 3.2.1 Enhanced brain activations for condition-specific contrasts

The main objective of fMRI data analysis was to focus on contrasting our conditions, the arousal value of which was entered as a parametric modulator on a trial-by-trial basis at the first-level of analysis. Importantly, because the aim of the present study was the investigation of emotional regulation for affective events and because the ratio between negative and neutral stimuli was strongly biased, only negative valence trials were included in the results presented here (see Methods, Neuroimaging data analysis). All activations are summarized in Table 1. Hence we first contrasted Watch against Reappraise trials and found a very specific cluster of enhanced activation in the right fusiform gyrus (Figure 3A; FFG, MNI xyz [38 -72 -9]). This cluster showed increased activity for Watch trials while Reappraise trials showed a decrease of activity (Figure 3B).

**Figure 3.**
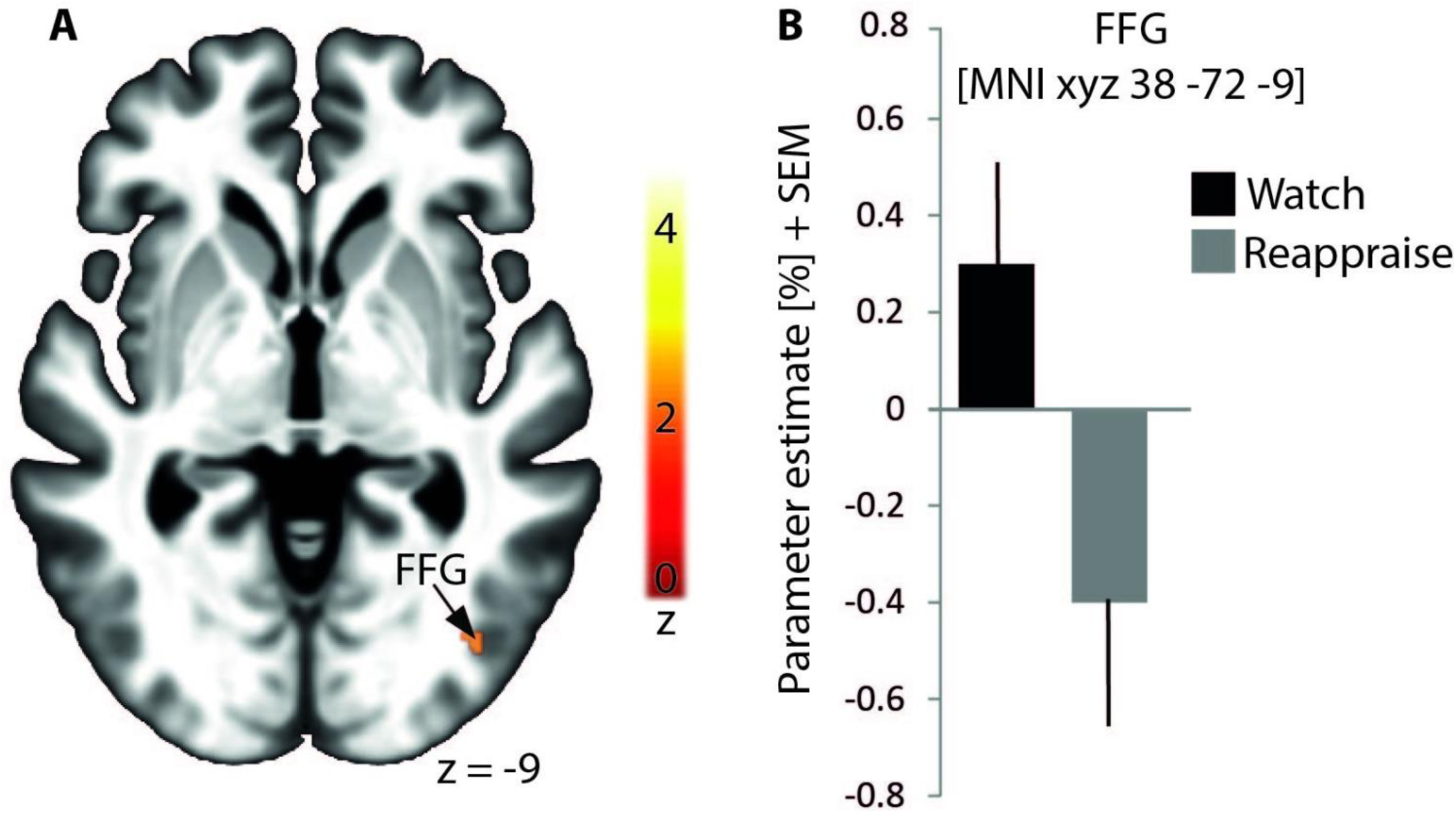
Watch > Reappraise. (A) Enhanced activity in the right fusiform gyrus (FFG; MNI xyz [38 −72 −9]). (B) Parameter estimates (± SEM) extracted at the peak voxel for Watch and Reappraise trials (negative images only), shown for illustration. Statistical maps are displayed at p < .005 uncorrected with a cluster extent threshold of k > 86 voxels (p < .05 FWE-corrected at the cluster level) on an age-specific template (Rorden et al., 2012).

**Table 1.**
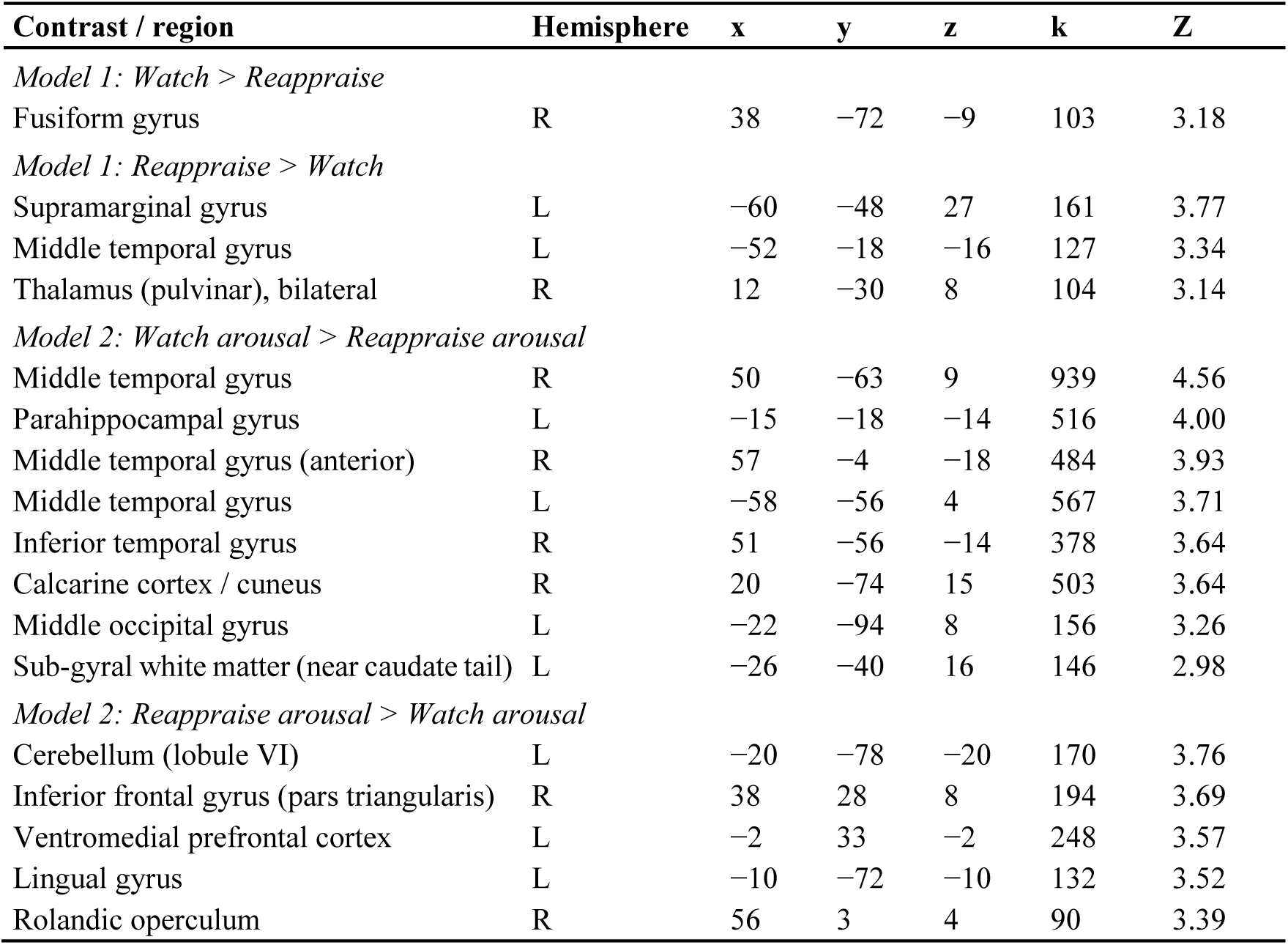
Peak activations for the condition-specific (Model 1, negative trials) and arousal-specific (Model 2) contrasts. Coordinates are in MNI space (mm); k, cluster size in voxels (1.5 × 1.5 × 1.5 mm); Z values were derived from the peak T values (df = 69). All clusters survived p < 0.005 uncorrected at the voxel level with a cluster extent threshold of k > 86 voxels (p < 0.05 FWE-corrected at the cluster level). Anatomical labels are based on the AAL atlas (Tzourio-Mazoyer et al., 2002). L, left; R, right.

We then contrasted Reappraise to Watch trials and found enhanced BOLD signal in the left supramarginal and middle temporal gyrus (Figure 4A; SMG, MNI xyz [-60 -48 27]; Figure 4B; MTG, MNI xyz [-51 -24 -11]). For both brain areas, signal extraction emphasized an increase of activity for Reappraise trials and a decrease of activity for Watch trials (Figure 4C).

**Figure 4.**
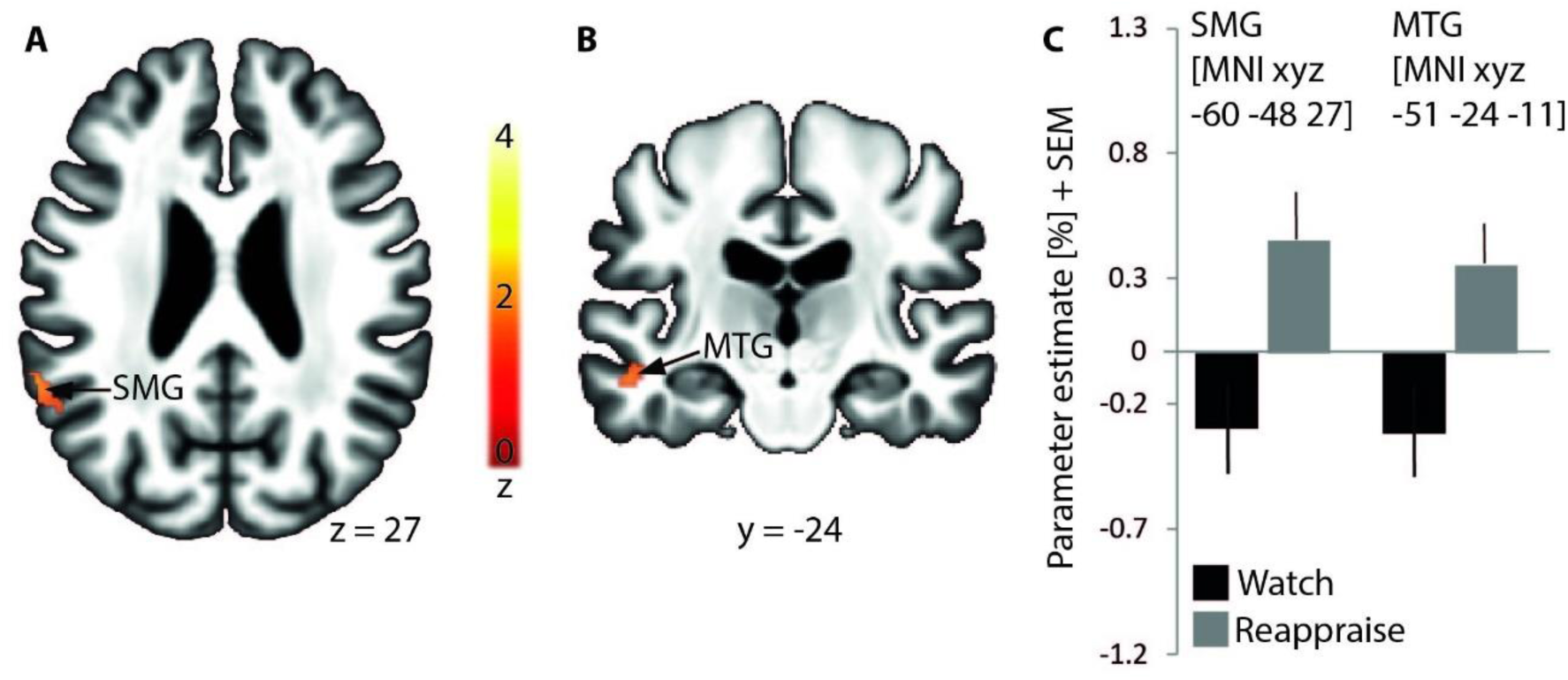
Reappraise > Watch. Enhanced activity in (A) the left supramarginal gyrus (SMG; MNI xyz [−60 −48 27]) and (B) the left middle temporal gyrus (MTG; MNI xyz [−51 −24 −11]). (C) Parameter estimates (± SEM) extracted at each peak for Watch and Reappraise trials (negative images only), shown for illustration. Thresholds and template as in Figure 3.

#### 3.2.2 Enhanced brain activations for arousal-specific contrasts of the main conditions

Following condition-specific analyses, we were interested in brain activity related to the arousing value of our negative stimuli, for each condition. We thus computed the Watch arousal > Reappraise arousal contrast and observed enhanced brain activations in the left parahippocampal gyrus (Figure 5A; PHG, MNI xyz [-15 -18 -14]), right middle temporal gyrus (Figure 5B; MTG, MNI xyz [50 -63 9]) and inferior temporal gyrus (Figure 5B; ITG, MNI xyz [51 -55 -14]). Since these second-level regressors involve values from a parametric modulator at the first-level, the results illustrate enhanced brain activity linearly increasing with the arousing value of Watch compared to Reappraise trials (Figure 5C). The bar plots therefore depict arousal slopes (parametric modulator estimates) rather than mean activity.

**Figure 5.**
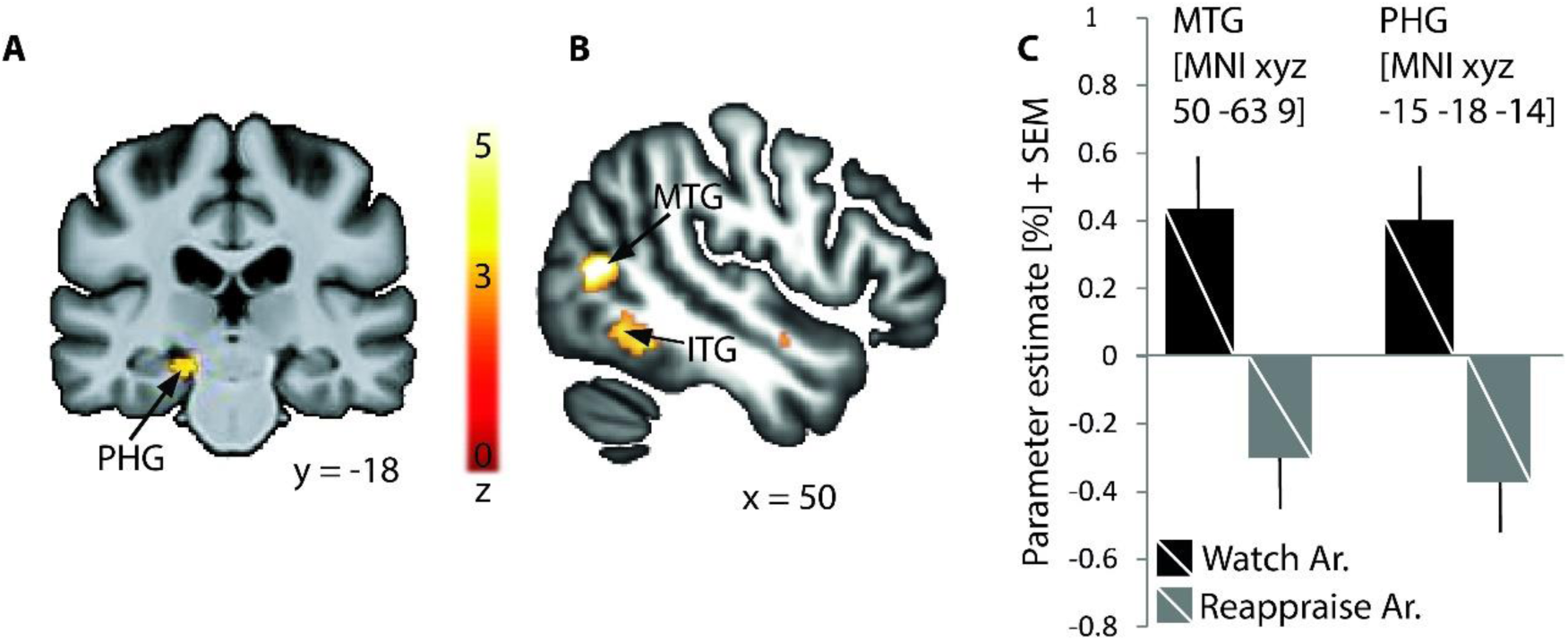
Watch arousal > Reappraise arousal. Steeper increase of activity with image arousal during watching than during reappraisal in (A) the left parahippocampal gyrus (PHG; MNI xyz [−15 −18 −14]) and (B) the right middle (MTG; MNI xyz [50 −63 9]) and inferior temporal gyri (ITG; MNI xyz [51 −55 −14]). (C) Arousal slopes (parametric modulator estimates ± SEM) extracted at the MTG and PHG peaks for Watch and Reappraise trials, shown for illustration. Thresholds and template as in Figure 3.

Contrasting Reappraise arousal to Watch arousal highlighted enhanced activity in the ventromedial prefrontal cortex (Figure 6A; vmPFC, MNI xyz [-2 33 -2]), the right inferior frontal gyrus (Figure 6B; IFG, MNI xyz [35 30 6]) and the right middle frontal gyrus (Figure 6B; MFG, MNI xyz [-54 15 6]). Once again, since these second-level regressors involve values from a parametric modulator at the first-level, the results illustrate enhanced brain activity linearly increasing with the arousing value of Reappraise compared to Watch trials (Figure 6C).

**Figure 6.**
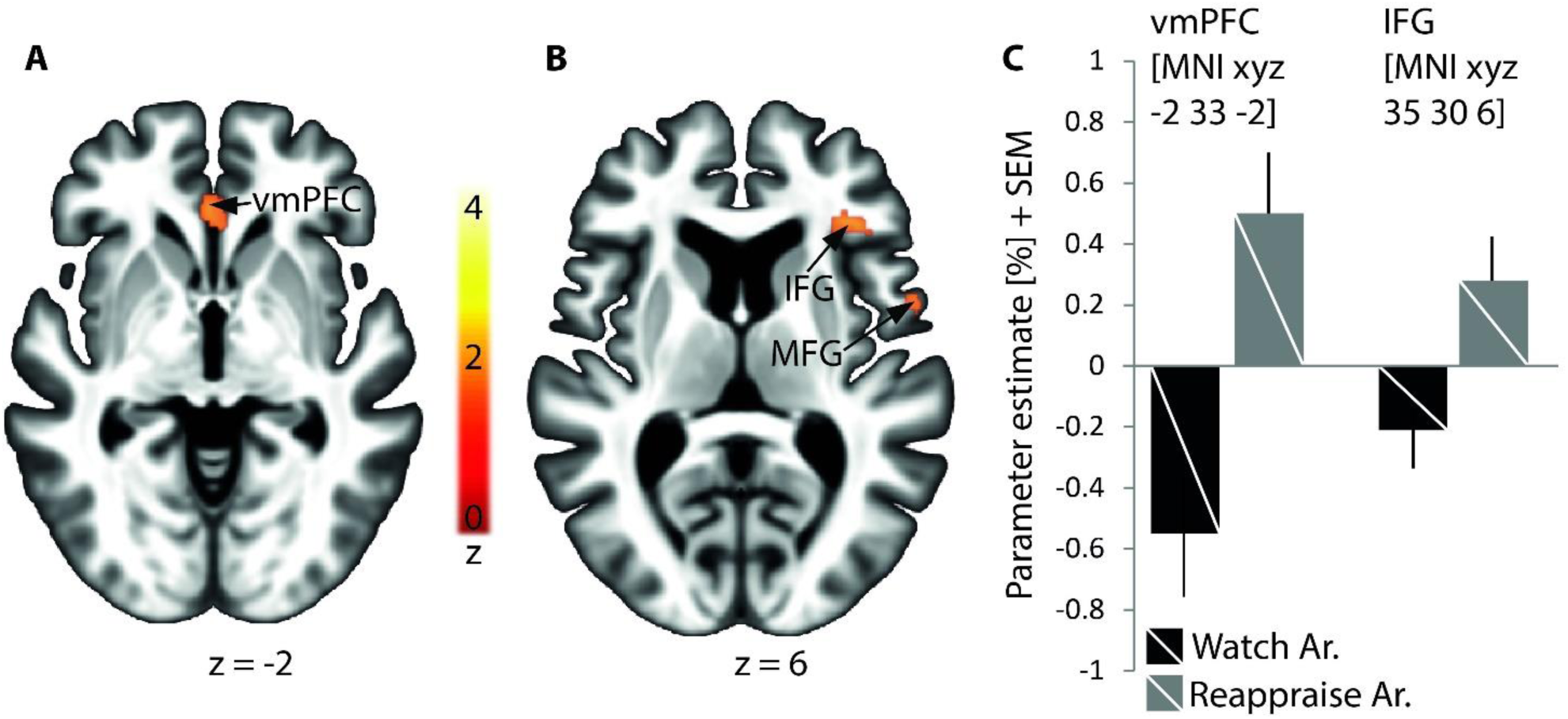
Reappraise arousal > Watch arousal. Steeper increase of activity with image arousal during reappraisal than during watching in (A) the ventromedial prefrontal cortex (vmPFC; MNI xyz [−2 33 −2]) and (B) the right inferior frontal gyrus (IFG; MNI xyz [35 30 6]) and middle frontal gyrus (MFG; MNI xyz [-54 15 6]). (C) Arousal slopes (parametric modulator estimates ± SEM) extracted at each peak for Watch and Reappraise trials, shown for illustration. Thresholds and template as in Figure 3.

#### 3.2.3 Seed-to-seed functional connectivity analyses

Contrasting Reappraise with Watch trials revealed condition differences in ten seed–target connections (p < 0.05, seed-level FDR-corrected; Figure 7; Table 2). All but one reflected stronger coupling during reappraisal than during watching. With the anterior cingulate cortex (AC) as seed, coupling increased with the right middle frontal gyrus (MidFG; T(23) = 4.51, p-FDR = 0.021), the right inferior frontal gyrus, pars triangularis (IFG tri; T(23) = 4.06, p-FDR = 0.031) and the left posterior supramarginal gyrus (pSMG; T(23) = 3.80, p-FDR = 0.040). With the posterior cingulate cortex (PC) as seed, coupling increased with the right MidFG (T(23) = 4.23, p-FDR = 0.035) and the right frontal pole (FP; T(23) = 4.02, p-FDR = 0.035).

**Figure 7.**
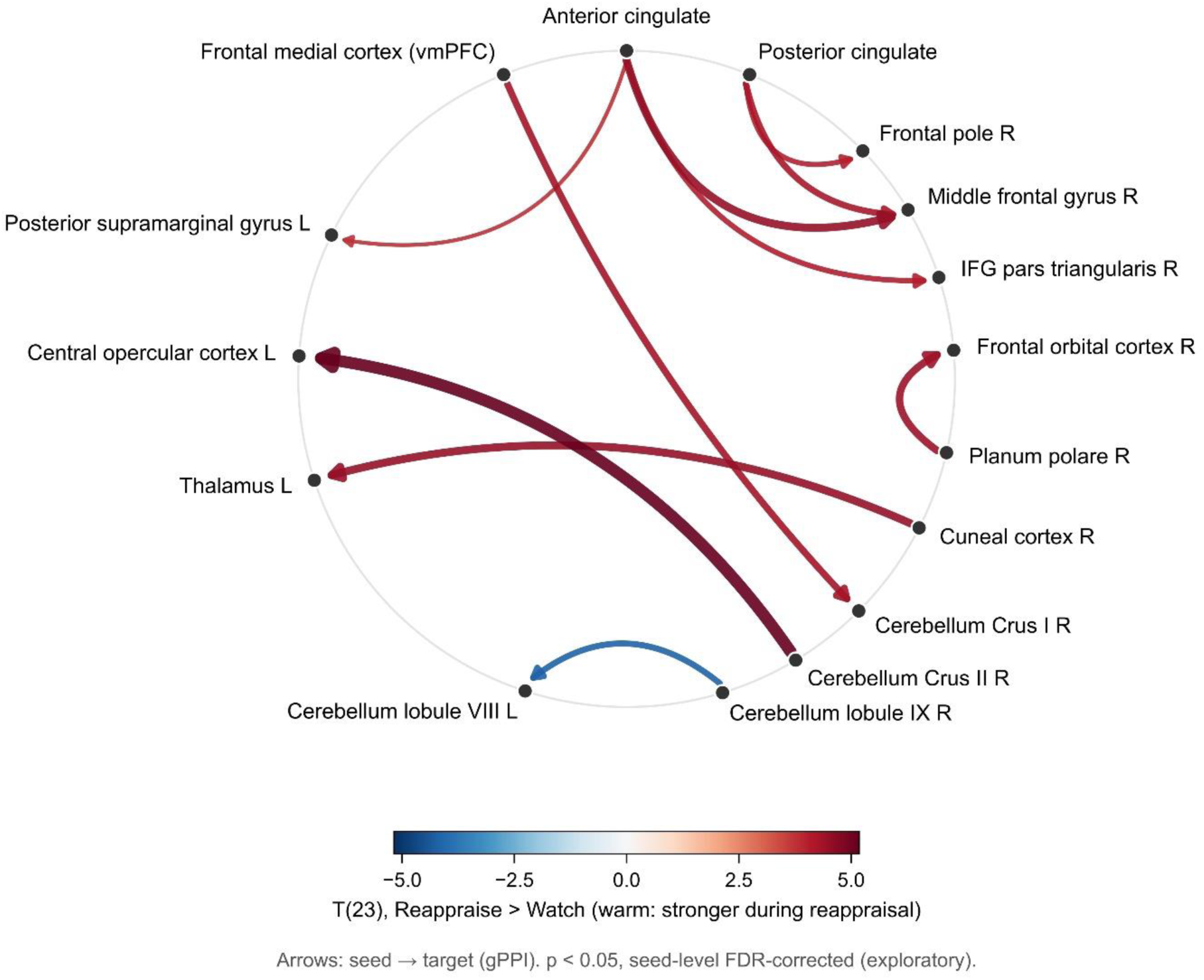
Exploratory seed-to-seed functional connectivity, Reappraise > Watch (gPPI; N = 24). Arrows point from seed to target ROI; arrow colour and width scale with the T value (warm colours: stronger coupling during reappraisal; cool colours: weaker coupling during reappraisal). Connections are displayed at p < .05, seed-level FDR-corrected. ROIs were taken from the CONN default atlas (Harvard-Oxford cortical and subcortical atlas, AAL cerebellar regions). IFG, inferior frontal gyrus; L, left; R, right; vmPFC, ventromedial prefrontal cortex.

**Table 2.**
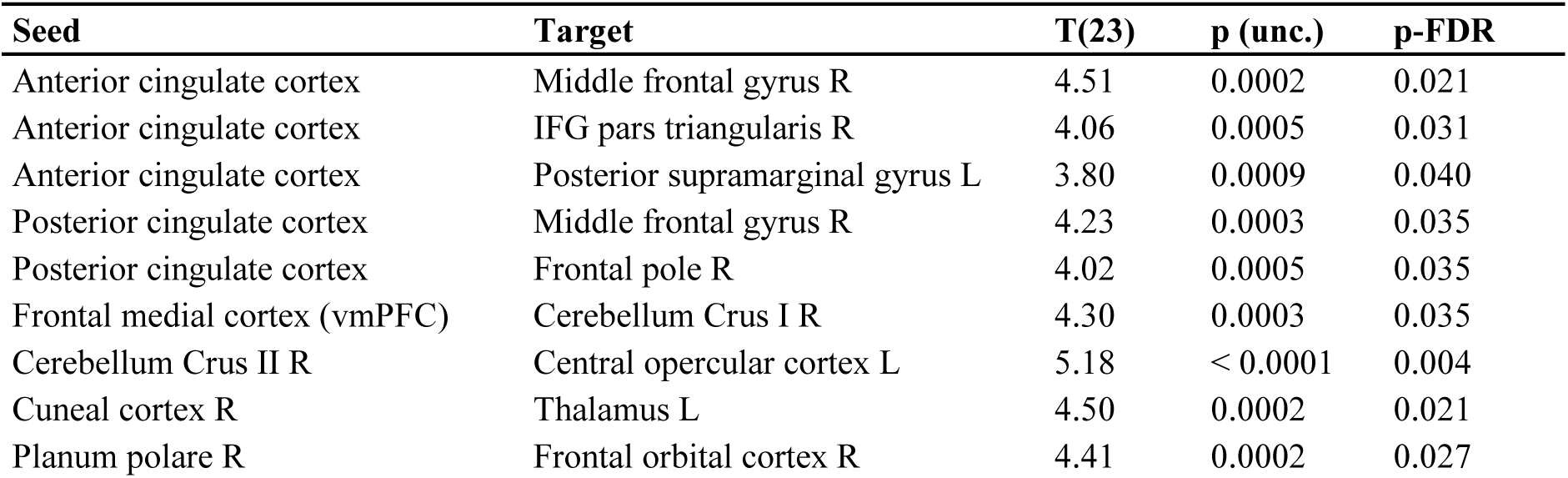
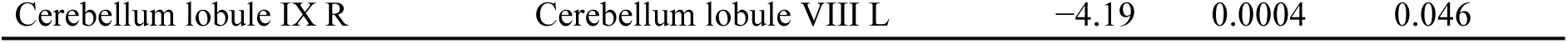
Exploratory seed-to-seed functional connectivity differences, Reappraise > Watch (gPPI, N = 24). Positive T values indicate stronger coupling during reappraisal than during watching. p-FDR, false discovery rate correction over the 131 connections of each seed (seed-level correction). IFG, inferior frontal gyrus; vmPFC, ventromedial prefrontal cortex; L, left; R, right.

Coupling during reappraisal also increased from the frontal medial cortex (MedFC, which encompasses the vmPFC) to right cerebellar Crus I (T(23) = 4.30, p-FDR = 0.035), and from right cerebellar Crus II to the left central opercular cortex (CO; T(23) = 5.18, p-FDR = 0.004), the strongest effect of the analysis. Further increases were found from the right cuneal cortex to the left thalamus (T(23) = 4.50, p-FDR = 0.021) and from the right planum polare (PP) to the right frontal orbital cortex (FOrb; T(23) = 4.41, p-FDR = 0.027). The only connection showing the reverse pattern was from right cerebellar lobule IX to left cerebellar lobule VIII, whose coupling decreased for Reappraise compared to Watch trials (T(23) = -4.19, p-FDR = 0.046). None of the seed-level omnibus tests survived FDR correction across seeds (all seed-level p-FDR > 0.20), so these connection-level effects should be considered exploratory.

## 4 Discussion

There is some evidence in the literature that older adults are more strongly motivated than young adults to maximize their current affective well-being (Carstensen, 2006). For instance Scheibe and colleagues found an age-related shift toward disengagement strategies (suppression) to regulate negative emotions, which maps onto older adults’ prohedonic orientation and holds affective benefits (Scheibe et al., 2015). Older adults could also rely on reappraisal in order to down-regulate their negative emotions. In the current study, we investigated reappraisal strategies and their neural substrate in a group of older adults. Behavioral analysis revealed that older participants were able to reappraise negative pictures as they rated the intensity of their negative emotions significantly lower in a reappraise condition compared to watch-only condition. At the neural level, our main finding is that arousal is a modulator on reappraised strategies in our group of older adults. Indeed, we discovered that the high arousal of reappraised images compared to watch images proportionally correlated with brain activity in the left ventromedial prefrontal cortex (vmPFC) and the right inferior frontal gyrus (IFG). Therefore, older adults might engage in a reappraise process only for highly aroused images.

In the context of aging, the interplay between emotion regulation and arousal is of particular theoretical interest because of concurrent age-related changes in physiological functioning and preferred affective experiences (Sands and Isaacowitz, 2017). Because older adults experience more difficulties to recover from heightened arousal, they might limit their exposure to potentially perturbing stimuli as predicted by the Socioemotional Selectivity Theory (Carstensen et al., 1999) and the Strength and Vulnerability Integration model (Charles, 2010). In that context, Wood et al. (2017) examined age differences in affect regulation following the dynamics-of-affect model (Kuppens et al., 2010) which estimated the attractor point (level of valence and arousal corresponding to the equilibrium state of an individual), reactivity to biopsychosocial inputs (valence and arousal variability associated with external and internal stimuli) and attractor strengths (intensity of the strength of bringing the individual to his initial attractor point). In-line with our results, they found that older age was associated with higher arousal attractor point and less BPS-related reactivity, but not with the dynamics of affect valence.

Going back to neuroimaging, the vmPFC comprises a set of interconnected regions that integrate information from affective sensory and social cues, long-term memory, and representations of the “self” (Roy et al., 2012). Over the past decade, neuroimaging studies have found the vmPFC to be a key region for various cognitive functions such as emotion regulation (Etkin et al., 2015), mentallization about others (Amodio and Frith, 2006) or even decision making (Hebscher and Gilboa, 2016). Despite the diverse nature of abovementioned functions, Roy et al. (2012) proposed that they all involve an affective responses that is shaped by conceptual information. Therefore, the vmPFC might be viewed as a hub, playing a critical role for the generation of affective responses driven by conceptual appraisals, a viewpoint that is highly compatible with the reappraisal condition of our task. Regarding the IFG, it has been proposed as a key player in reconfiguring a representation of the currently attended input (Hampshire et al., 2010). Indeed, the IFG has been classically associated with inhibitory control (Aron et al., 2004) or more recently as a brake over a strong response tendency (Aron et al., 2014). Coherent with these findings, the IFG might be, in the context of our task, more strongly activated when a highly aroused image has to be reappraised. Additional clusters are listed in Table 1.

At the network level, exploratory functional connectivity analyses complemented these activation results. Reappraising rather than watching negative pictures was associated with stronger coupling between the anterior cingulate cortex and the right inferior and middle frontal gyri, and between the posterior cingulate cortex and right lateral prefrontal regions (middle frontal gyrus, frontal pole). This pattern fits models in which reappraisal relies on interactions between cingulate and lateral prefrontal control regions (Ochsner et al., 2012; Kohn et al., 2014), and suggests that the right IFG highlighted by our arousal-related analysis may operate within a broader cingulo-prefrontal circuit during reappraisal. The anterior cingulate cortex was also more strongly coupled with the left posterior supramarginal gyrus, that is, the gyrus in which we also observed enhanced activity for Reappraise compared to Watch trials, thereby linking our activation and connectivity findings. In addition, the medial frontal cortex, which includes the vmPFC, was more strongly coupled with right cerebellar Crus I during reappraisal. Crus I and Crus II belong to the cerebellar territories that are functionally coupled with prefrontal association cortices (Buckner et al., 2011), and cerebellar lesions can impair executive and affective functions (Schmahmann and Sherman, 1998). Our results therefore point to a possible cerebellar contribution to reappraisal in older adults, a hypothesis that remains to be tested directly. These connectivity findings should nevertheless be interpreted with caution: multiple comparisons were corrected within each seed rather than across the whole network, and no seed-level omnibus test survived correction.

Contrary to previous studies exploring emotion regulation in young (e.g., Kohn et al., 2014) or older adults (van Reekum et al., 2007; Urry et al., 2009), we did not observed that reappraise lead to an increase activation of dorsolateral prefrontal cortex (dlPFC) and dorsomedial prefrontal cortex (dmPFC), along with the inferior parietal lobule (IPL). Interestingly, Urry and colleagues (2006) observed that subjects showing the greatest reappraisal-related decrease in amygdala activation also showed greater activation in the vmPFC but not in the more typically reported medial and lateral regions. The authors concluded that the VMPFC was implicated in the ability of subjects to transform their experience of negative stimuli to be less negative (Urry et al., 2006).

There are nevertheless some limitations that need to be addressed in our study. First, older adults might have deployed attention away from negative pictures as some studies found they preferred using suppression strategies to downregulate negative emotions (Scheibe et al., 2015). The use of eye-tracking methodology would have been useful to determine whether older adults try to avoid focusing their attention on negative pictures or highly salient/unpleasant areas of the images. While we tried to collect eye tracking data during our experiment, technical difficulties didn’t allow us to use the data (recording problems). A direct assessment of visual attention should hence be addressed in future experiments. Secondly, it would be important to have a more objective measure of the effect of reappraisal by collecting physiological data. For instance, skin conductance response could provide a measure of sympathetic nervous system activity, associated with phasic increases in emotional arousal (Demaree et al., 2004).

Our study highlights that arousal plays an important role in older adults as reappraisal strategies seem to be modulated by this emotional dimension, according to our data. The arousal deficit hypothesis suggests that « positive bias » in older adults might come from a change in the early perceptual processing (Corbetta and Shulman, 2002, at the earlier arousal stage, « ventral attention »; Bressler et al., 2008) rather than at a latter top-down cognitive processing stage (« dorsal attention »). It would be important in the future to explore more deeply the bottom-up treatment of negatively valenced visual stimuli in older adults, notably by enhancing the arousal dimension using for instance dynamic rather than static visual stimuli.

Some limitations of the present study should be acknowledged. First, the absence of a younger control group prevents us from determining whether the observed behavioral and neural effects are specific to older adults or reflect age-invariant mechanisms of reappraisal; age-related interpretations should therefore be considered tentative and tested in future studies directly comparing age groups. Second, activations were thresholded at p < 0.005 uncorrected at the voxel level combined with a cluster-extent correction; such cluster-defining thresholds have been associated with inflated false-positive rates (Eklund et al., 2016), and the present activation results would therefore benefit from replication in an independent sample. Third, captions suggesting a possible reappraisal (Reappraise trials) or describing the scene (Watch trials) were displayed with the pictures in the second run, so that reappraisal was partly guided in this run, whereas ratings and brain responses were pooled across runs. Finally, the functional connectivity results were corrected for multiple comparisons within each seed only and should be considered exploratory.

## 5 Data Availability Statement

All data and codes of this article are freely available on the FAIR-compliant, CoreTrustSeal-certified Yareta repository at: https://doi.org/10.26037/yareta:axwpfp2lz5h2zgmqehkvupvgwa.

## 6 Ethics Statement

The studies involving human participants were reviewed and approved by the cantonal research ethics committee of Geneva (Commission Cantonale d’Éthique de la Recherche, CCER). The participants provided their written informed consent to participate in this study.

## 8 Author Contributions

LC acquired the behavioral and imaging data, analyzed them and wrote the manuscript. GM acquired part of the behavioral, MRI data, was in charge of the ‘GDS’, ‘Mini’ and ‘Mattis’ evaluations (inclusion and recruitment) and took part in manuscript writing. CS took part in the inclusion and recruitment process. AV acquired the behavioral data, part of the MRI data, and took part in the inclusion and recruitment process. DF took part in the inclusion and recruitment process and in the neuropsychological evaluations. GG, FL and PG helped improve the manuscript. PV helped analyze MRI data and improve the manuscript. CD managed the whole team and took part in inclusion and recruitment, behavioral data analysis and manuscript editing and writing. All authors took part in manuscript writing and/or editing.

## 10 Acknowledgments

The authors would like to thank Bruno Bonet and Frédéric Grouiller for their great help regarding neuroimaging data acquisition. The authors also warmly thank our participants for agreeing to take part in this study.

## References

1. Amodio, D. M., and Frith, C. D. (2006). Meeting of minds: the medial frontal cortex and social cognition. Nat. Rev. Neurosci. 7, 268–277.

2. Aron, A. R., Robbins, T. W., and Poldrack, R. A. (2004). Inhibition and the right inferior frontal cortex. Trends Cogn. Sci. 8, 170–177.

3. Aron, A. R., Robbins, T. W., and Poldrack, R. A. (2014). Inhibition and the right inferior frontal cortex: one decade on. Trends Cogn. Sci. 18, 177–185.

4. Ashburner, J. (2007). A fast diffeomorphic image registration algorithm. Neuroimage 38, 95–113.

5. Behzadi, Y., Restom, K., Liau, J., and Liu, T. T. (2007). A component based noise correction method (CompCor) for BOLD and perfusion based fMRI. Neuroimage 37, 90–101.

6. Benjamini, Y., and Hochberg, Y. (1995). Controlling the false discovery rate: a practical and powerful approach to multiple testing. J. R. Stat. Soc. Series B Stat. Methodol. 57, 289–300.

7. Blazer, D. G. (2003). Depression in late life: review and commentary. J. Gerontol. A Biol. Sci. Med. Sci. 58, M249–M265.

8. Bressler, S. L., Tang, W., Sylvester, C. M., Shulman, G. L., and Corbetta, M. (2008). Top-down control of human visual cortex by frontal and parietal cortex in anticipatory visual spatial attention. J. Neurosci. 28, 10056–10061.

9. Buckner, R. L., Krienen, F. M., Castellanos, A., Diaz, J. C., and Yeo, B. T. (2011). The organization of the human cerebellum estimated by intrinsic functional connectivity. J. Neurophysiol. 106, 2322– 2345.

10. Carstensen, L. L. (2006). The influence of a sense of time on human development. Science 312, 1913–1915.

11. Carstensen, L. L., Isaacowitz, D. M., and Charles, S. T. (1999). Taking time seriously: a theory of socioemotional selectivity. Am. Psychol. 54, 165–181.

12. Charles, S. T. (2010). Strength and vulnerability integration: a model of emotional well-being across adulthood. Psychol. Bull. 136, 1068–1091.

13. Collins, D. L., Neelin, P., Peters, T. M., and Evans, A. C. (1994). Automatic 3D intersubject registration of MR volumetric data in standardized Talairach space. J. Comput. Assist. Tomogr. 18, 192–205.

14. Corbetta, M., and Shulman, G. L. (2002). Control of goal-directed and stimulus-driven attention in the brain. Nat. Rev. Neurosci. 3, 201–215.

15. Cox, R. W. (1996). AFNI: software for analysis and visualization of functional magnetic resonance neuroimages. Comput. Biomed. Res. 29, 162–173.

16. Demaree, H. A., Schmeichel, B. J., Robinson, J. L., and Everhart, D. E. (2004). Behavioural, affective, and physiological effects of negative and positive emotional exaggeration. Cogn. Emot. 18, 1079–1097.

17. Desikan, R. S., Ségonne, F., Fischl, B., Quinn, B. T., Dickerson, B. C., Blacker, D., et al. (2006). An automated labeling system for subdividing the human cerebral cortex on MRI scans into gyral based regions of interest. Neuroimage 31, 968–980.

18. Desmond, J. E., and Glover, G. H. (2002). Estimating sample size in functional MRI (fMRI) neuroimaging studies: statistical power analyses. J. Neurosci. Methods 118, 115–128.

19. Ebert, D., and Ebmeier, K. P. (1996). The role of the cingulate gyrus in depression: from functional anatomy to neurochemistry. Biol. Psychiatry 39, 1044–1050.

20. Ehring, T., Tuschen-Caffier, B., Schnülle, J., Fischer, S., and Gross, J. J. (2010). Emotion regulation and vulnerability to depression: spontaneous versus instructed use of emotion suppression and reappraisal. Emotion 10, 563–572.

21. Eklund, A., Nichols, T. E., and Knutsson, H. (2016). Cluster failure: why fMRI inferences for spatial extent have inflated false-positive rates. Proc. Natl. Acad. Sci. U.S.A. 113, 7900–7905.

22. Erk, S., Mikschl, A., Stier, S., Ciaramidaro, A., Gapp, V., Weber, B., et al. (2010). Acute and sustained effects of cognitive emotion regulation in major depression. J. Neurosci. 30, 15726–15734.

23. Etkin, A., Büchel, C., and Gross, J. J. (2015). The neural bases of emotion regulation. Nat. Rev. Neurosci. 16, 693–700.

24. Faul, F., Erdfelder, E., Lang, A.-G., and Buchner, A. (2007). G*Power 3: a flexible statistical power analysis program for the social, behavioral, and biomedical sciences. Behav. Res. Methods 39, 175– 191.

25. Frazier, J. A., Chiu, S., Breeze, J. L., Makris, N., Lange, N., Kennedy, D. N., et al. (2005). Structural brain magnetic resonance imaging of limbic and thalamic volumes in pediatric bipolar disorder. Am. J. Psychiatry 162, 1256–1265.

26. Goldstein, J. M., Seidman, L. J., Makris, N., Ahern, T., O’Brien, L. M., Caviness, V. S., et al. (2007). Hypothalamic abnormalities in schizophrenia: sex effects and genetic vulnerability. Biol. Psychiatry 61, 935–945.

27. Gross, J. J. (1998). The emerging field of emotion regulation: an integrative review. Rev. Gen. Psychol. 2, 271–299.

28. Gross, J. J. (1999). Emotion regulation: past, present, future. Cogn. Emot. 13, 551–573.

29. Gross, J. J., and Muñoz, R. F. (1995). Emotion regulation and mental health. Clin. Psychol. Sci. Pract. 2, 151–164.

30. Hampshire, A., Chamberlain, S. R., Monti, M. M., Duncan, J., and Owen, A. M. (2010). The role of the right inferior frontal gyrus: inhibition and attentional control. Neuroimage 50, 1313–1319.

31. Hebscher, M., and Gilboa, A. (2016). A boost of confidence: the role of the ventromedial prefrontal cortex in memory, decision-making, and schemas. Neuropsychologia 90, 46–58.

32. Johnstone, T., van Reekum, C. M., Urry, H. L., Kalin, N. H., and Davidson, R. J. (2007). Failure to regulate: counterproductive recruitment of top-down prefrontal-subcortical circuitry in major depression. J. Neurosci. 27, 8877–8884.

33. Kohn, N., Eickhoff, S. B., Scheller, M., Laird, A. R., Fox, P. T., and Habel, U. (2014). Neural network of cognitive emotion regulation—an ALE meta-analysis and MACM analysis. Neuroimage 87, 345– 355.

34. Kuppens, P., Oravecz, Z., and Tuerlinckx, F. (2010). Feelings change: accounting for individual differences in the temporal dynamics of affect. J. Pers. Soc. Psychol. 99, 1042–1060.

35. Lang, P. J., Bradley, M. M., and Cuthbert, B. N. (2008). International Affective Picture System (IAPS): Affective Ratings of Pictures and Instruction Manual. Technical Report A-8. Gainesville, FL: University of Florida.

36. Lau, A. W., Edelstein, B. A., and Larkin, K. T. (2001). Psychophysiological arousal in older adults: a critical review. Clin. Psychol. Rev. 21, 609–630.

37. Lecrubier, Y., Sheehan, D. V., Weiller, E., Amorim, P., Bonora, I., Sheehan, K. H., et al. (1997). The Mini International Neuropsychiatric Interview (MINI). A short diagnostic structured interview: reliability and validity according to the CIDI. Eur. Psychiatry 12, 224–231.

38. Makris, N., Goldstein, J. M., Kennedy, D., Hodge, S. M., Caviness, V. S., Faraone, S. V., et al. (2006). Decreased volume of left and total anterior insular lobule in schizophrenia. Schizophr. Res. 83, 155– 171.

39. Mattis, S. (1976). “Mental status examination for organic mental syndrome in the elderly patient,” in Geriatric Psychiatry, eds L. Bellak and T. B. Karasu (New York, NY: Grune & Stratton), 77–121.

40. McLaren, D. G., Ries, M. L., Xu, G., and Johnson, S. C. (2012). A generalized form of context-dependent psychophysiological interactions (gPPI): a comparison to standard approaches. Neuroimage 61, 1277–1286.

41. Ochsner, K. N., Silvers, J. A., and Buhle, J. T. (2012). Functional imaging studies of emotion regulation: a synthetic review and evolving model of the cognitive control of emotion. Ann. N. Y. Acad. Sci. 1251, E1–E24.

42. Rorden, C., Bonilha, L., Fridriksson, J., Bender, B., and Karnath, H.-O. (2012). Age-specific CT and MRI templates for spatial normalization. Neuroimage 61, 957–965.

43. Roy, M., Shohamy, D., and Wager, T. D. (2012). Ventromedial prefrontal-subcortical systems and the generation of affective meaning. Trends Cogn. Sci. 16, 147–156.

44. Sands, M., and Isaacowitz, D. M. (2017). Situation selection across adulthood: the role of arousal. Cogn. Emot. 31, 791–798.

45. Scheibe, S., Sheppes, G., and Staudinger, U. M. (2015). Distract or reappraise? Age-related differences in emotion-regulation choice. Emotion 15, 677–681.

46. Scherer, K. R. (1984). Emotion as a multicomponent process: a model and some cross-cultural data. Rev. Pers. Soc. Psychol. 5, 37–63.

47. Schmahmann, J. D., and Sherman, J. C. (1998). The cerebellar cognitive affective syndrome. Brain 121, 561–579.

48. Thiruchselvam, R., Blechert, J., Sheppes, G., Rydstrom, A., and Gross, J. J. (2011). The temporal dynamics of emotion regulation: an EEG study of distraction and reappraisal. Biol. Psychol. 87, 84– 92.

49. Tzourio-Mazoyer, N., Landeau, B., Papathanassiou, D., Crivello, F., Etard, O., Delcroix, N., et al. (2002). Automated anatomical labeling of activations in SPM using a macroscopic anatomical parcellation of the MNI MRI single-subject brain. Neuroimage 15, 273–289.

50. Urry, H. L., van Reekum, C. M., Johnstone, T., and Davidson, R. J. (2009). Individual differences in some (but not all) medial prefrontal regions reflect cognitive demand while regulating unpleasant emotion. Neuroimage 47, 852–863.

51. Urry, H. L., van Reekum, C. M., Johnstone, T., Kalin, N. H., Thurow, M. E., Schaefer, H. S., et al. (2006). Amygdala and ventromedial prefrontal cortex are inversely coupled during regulation of negative affect and predict the diurnal pattern of cortisol secretion among older adults. J. Neurosci. 26, 4415–4425.

52. van Reekum, C. M., Johnstone, T., Urry, H. L., Thurow, M. E., Schaefer, H. S., Alexander, A. L., et al. (2007). Gaze fixations predict brain activation during the voluntary regulation of picture-induced negative affect. Neuroimage 36, 1041–1055.

53. Wager, T. D., Davidson, M. L., Hughes, B. L., Lindquist, M. A., and Ochsner, K. N. (2008). Prefrontal-subcortical pathways mediating successful emotion regulation. Neuron 59, 1037–1050.

54. Whitfield-Gabrieli, S., and Nieto-Castanon, A. (2012). Conn: a functional connectivity toolbox for correlated and anticorrelated brain networks. Brain Connect. 2, 125–141.

55. Wood, J., Oravecz, Z., Vogel, N., Benson, L., Chow, S.-M., Cole, P., et al. (2017). Modeling intraindividual dynamics using stochastic differential equations: age differences in affect regulation. J. Gerontol. B Psychol. Sci. Soc. Sci. doi: 10.1093/geronb/gbx013

56. Wood, S., and Kisley, M. A. (2006). The negativity bias is eliminated in older adults: age-related reduction in event-related brain potentials associated with evaluative categorization. Psychol. Aging 21, 815–820.

57. Yesavage, J. A., Brink, T. L., Rose, T. L., Lum, O., Huang, V., Adey, M., et al. (1983). Development and validation of a geriatric depression screening scale: a preliminary report. J. Psychiatr. Res. 17, 37– 49.

